# Astrocytes control the Neuroinflammation and ILC2 response through IL-33/ST2 signaling, during protection against Cerebral Malaria in *Toxoplasma*-*P. berghei* coinfected Mice

**DOI:** 10.1101/2025.10.24.684259

**Authors:** Ines Leleu, Tarun Keswani, Fabien Herbert, Capucine Picavet, Manon Arnaud, Marion Sabrina, Corine Glineur, Pied Sylviane

## Abstract

Cerebral malaria (CM) is a complex multi-systemic disorder defined as a diffuse encephalopathy with acute neurological manifestations characterized by alterations in the level of consciousness, deep coma and seizure preceding death. During infection, astrocytes undergo significant morphological and molecular changes, adopting a reactive state that impacts on their functions. This reactivity is characterized by a shift in from a neuroprotective (A2) to a neurotoxic (A1) phenotype, influencing the outcome of the immune response. These phenotypes may vary depending on the chronicity of the infection or multiples infections of the same host.

In this study, we investigated how *Toxoplasma gondii* (*Tg*) brain infection impacts on the outcome of experimental cerebral malaria (ECM) in mice infected with *Plasmodium berghei* ANKA (*Pb*A). Our results highlighted an immunomodulatory role of GFAP^+^ astrocytes underweening significant morphological and molecular alterations and adopting a unique intermediate reactivity state (A1/A2). This state was correlated with production of CXCL-10 and TGF-β, which control inflammation without exacerbating infection. Our study also revealed a key role of the IL-33/ST2 pathway induced by *Tg* brain infection in protecting against ECM. Astrocyte-derived IL-33 was crucial to promote brain recruitment and activation of innate lymphoid cells (ILC2), which contribute to the host’s antiparasitic response. Additionally, we identified a distinctive intermediate M1/M2 phenotype in CD86^+^CD206^+^CD16/32^+^MHCII^hi^ microglia and noted an enhanced recruitment of inflammatory monocytes, both contributing to inflammation and control of *Pb*A infection.

This study reveals, for the first time, how latent brain infection with *T. gondii* confers protection against a severe cerebral form of malaria, positioning astrocytes at the core of the neuroinflammatory response that controls *Pb*A infection severity. This expands our understanding of host-pathogen interactions and the potential for targeting astrocytic pathways in preventing CM.

**Author Summary:** Cerebral malaria (CM) is one of the most severe complications of *Plasmodium* infection, often leading to coma and death. The mechanisms that determine why some individuals develop this life-threatening condition remain poorly understood. In this study, we explored how a chronic brain infection with the parasite *Toxoplasma gondii* influences the development of CM in mice. We found that *Tg* infection reshapes the brain’s immune environment, particularly through the actions of astrocytes, cells that normally support and protect neurons. During coinfection, astrocytes adopted a balanced reactive state that limited inflammation without worsening the infection. This response involved the IL-33/ST2 signalling pathway and led to the recruitment of protective immune cells, helping to control *Plasmodium* infection in the brain. Our findings uncover an unexpected protective role of latent *T. gondii* infection and identify astrocytes as central regulators of neuroinflammation. This work highlights potential new strategies for preventing or mitigating cerebral malaria by targeting astrocyte-mediated immune responses.

## Introduction

Coinfections with protozoan parasites targeting the central nervous system (CNS) are common in endemic regions, yet remain poorly understood (1–3). Once considered immune-privileged, the CNS is now recognized as highly vulnerable to immune-driven disruptions during infection. Neuroparasitic diseases such as *Plasmodium falciparum* cerebral malaria (CM) and *Toxoplasma gondii* (*Tg*) toxoplasmosis are among the most devastating, often leaving survivors with lifelong neurological sequelae (4–6). Even underreported, epidemiological data suggest a high prevalence of *Tg* in most malaria endemic areas (5,7,8). Despite their prevalence, neuroparasitic coinfections remain a neglected area of research, even though they can drastically modify host immunity and clinical outcomes.

CM, he most severe complication of *Plasmodium falciparum* infection, causes nearly 0.6 million deaths annually and is characterized by acute encephalopathy with coma and seizures(5). It is a complex multi-systemic disorder results from the combination of vascular obstruction by infected red blood cells (iRBCs) and a dysregulated neuroinflammatory response (Baptista et al. 2010). The experimental model of CM (ECM), induced by *Plasmodium berghei* ANKA (*Pb*A) in C57BL/6 mice, has been essential to dissect mechanisms of neuropathology. A central feature of ECM is the exacerbated activation of glial cells and the production of pro-inflammatory mediators such as CXCL-10 (C-X-C motif chemokine ligand-10), CCL-2 (C-C motif chemokine ligand-2), TNF-α (tumour necrosis factor), and interferon gamma (IFN-γ), which promote the selective recruitment of pathogenic αβCD8^+^ T cells to the brain (9–15). These T cells, by producing cytotoxic molecules (granzyme B, perforin) and inflammatory cytokines, disrupt the blood–brain barrier (BBB) and drive acute encephalopathy (14,16,17).

Astrocytes have emerged as central players in this pathological cascade. As the most abundant glial cells in the CNS, astrocytes normally contribute to tissue homeostasis and infection control (18). However, in ECM, they undergo profound morphological and functional changes and adopt a predominantly A1 neurotoxic phenotype, amplifying pro-inflammatory responses (9,10,19). Astrocytes are now recognized as a highly plastic population that can polarize into distinct functional phenotypes: the A1 subtype, characterized by a pro-inflammatory and neurotoxic profile, and the A2 subtype, associated with anti-inflammatory, neuroprotective, and tissue repair functions (19). Recent work demonstrated that *Pb*A-derived microvesicles are key drivers of this reprogramming: they are internalized by astrocytes through a Rubicon-dependent non-canonical autophagy pathway, leading to astrocyte senescence (4,12). Senescent astrocytes release large amounts of CXCL-10 and other chemokines, thereby probably fuelling the massive recruitment of CXCR3^+^ (C-X-C Motif Chemokine Receptor 3) CD8^+^ T cells into the brain (4,12,15,20). This creates a feed-forward loop of neuroinflammation, BBB breakdown, and neuronal dysfunction, hallmarks of ECM pathology (19,21). Thus, astrocyte dysfunction is not merely a bystander effect but a driving force of ECM immunopathology.

In contrast to ECM, *Tg* infection drives a more balanced and often protective astrocytic response (22–24). During chronic toxoplasmosis, astrocytes not only limit parasite dissemination by producing CXCL-10, CCL-2, and interleukin-6 (IL-6), but also promote antigen presentation to CD8^+^ T cells, thereby sustaining parasite control (25–29). Importantly, astrocytes in this context often adopt an A2-like phenotype, characterized by anti-inflammatory and neuroprotective properties. They contribute to the establishment of chronic infection rather than acute pathology by secreting TGF-β (Transforming Growth Factor-beta), which dampens excessive inflammation, preserves tissue integrity, and promotes tolerance to the presence of cysts (30). In addition, astrocytes release IL-33 as an alarmin that signals through the ST2 (Suppression of Tumorigenicity 2) receptor, orchestrating the recruitment of protective immune populations such as the Group 2 Innate Lymphoid Cells (ILC2s), IFN-γ–producing T cells, and iNOS^+^ (inducible Nitric Oxide Synthase) monocytes (31,32). Thus, in toxoplasmosis, astrocytes act not only as sentinels but also as immunoregulatory hubs that prevent uncontrolled inflammation and support long-term brain resistance.

Because *Plasmodium* and *Toxoplasma* overlap geographically and share CNS tropism, their coinfection raises crucial questions. Previous findings suggested that chronic *Tg* infection or *Tg* antigen exposure can attenuate ECM severity, pointing toward astrocyte-mediated immunoregulation as a protective mechanism (33). In this study, using a model of coinfection with *Tg* type II Prugniaud-green fluorescent protein (GFP) and *Pb*A in C57BL/6 mice, we explored how astrocytes integrate signals from both parasites to shape neuroinflammation. We demonstrate that protection against ECM depends on *Tg* cyst development and is not linked to parasite clearance but rather to astrocyte-driven immune modulation. In particular, our results highlight the importance of astrocyte polarization and signalling through the IL-33/ST2 pathway, which recruits ILC2s and balances inflammatory responses, thereby preventing fatal immunopathology.

## Materiel and Methods

### Mice

Female C57BL/6JRj (6–8 weeks, Janvier Laboratories, France) and IL-33 knockout (IL-33 KO) mice (34) were housed under specific pathogen-free conditions at the Institut Pasteur de Lille. Experiments were performed in accordance with institutional, French (Decree 87-848), European (Directive 86/609/CEE), and NIH (A5476-01) guidelines, with protocols approved by the Ethical Committee of Institute Pasteur of Lille and the French Committee for Animal Health Care “Ministère de l’Agriculture et de la Pêche” (#22021-2019091715283871).

### Toxoplasma gondii infection

Type II *T. gondii* tachyzoites (Prugniaud Pru-Luc-GFP strain) were maintained in human foreskin fibroblasts cultured in Dulbecco’s Modified Eagle Medium (DMEM; Gibco; 11.885-084) supplemented with 10% fetal bovine serum (FBS; Sigma-Aldrich; F0926), glutamine (Gibco; 25030081), and penicillin/streptomycin (P/S; Gibco; 10378016). Mice were infected intraperitoneally with 1×10³ tachyzoites.

### T. gondii parasite burden and cyst quantification

Genomic DNA (gDNA) was extracted from brain homogenate using the NucleoSpin Tissue DNA extraction kit (Macherey-Nagel; 740952.5) according to manufacturer’s recommendations. gDNA was quantified using a Nanodrop and the number of parasites per µg of brain DNA was calculated by real-time qPCR. Briefly, the *T. gondii* B1 gene was amplified using SYBR GreenER SuperMix for iCycler (Invitrogen; 11761500) with an MgCl_2_ concentration adjusted to 3.5µM in a 30µl reaction volume, 1 µg of total template DNA from each organ, 1.50µl of 0.5µM primer (sequences referencing in Supplementary Table 1 and water for a final volume of 30µl. Gene amplification was done with an iCycler^TM^ reverse transcription (RT)-PCR (Polymerase Chain Reaction) machine (Bio-Rad Laboratories) using a 10min initial denaturation at 95°C, followed by 50 cycles that consisted of 15s of denaturation at 95°C, 30s of annealing at 60°C, and 30s of extension at 72°C. A standard curve was used to made of a known number of corresponding parasites. Cysts were enumerated from brain homogenates labelled with FITC (Fluorescein Isothiocyanate)-conjugated Dolichos biflorus agglutinin (Vector Laboratories; FL-1031) and visualized by fluorescence microscopy with a 20X Objective.

### Experimental Cerebral malaria (ECM) model

ECM was induced by intraperitoneal infection with 1×10⁶ *Pb*A-iRBCs (ECM^+^; *Pb*A group in Figure 1A) (11). Parasitemia was assessed on Giemsa-stained blood smears at 6.5 days post-infection, when ECM symptoms typically developed.

**Figure 1.**
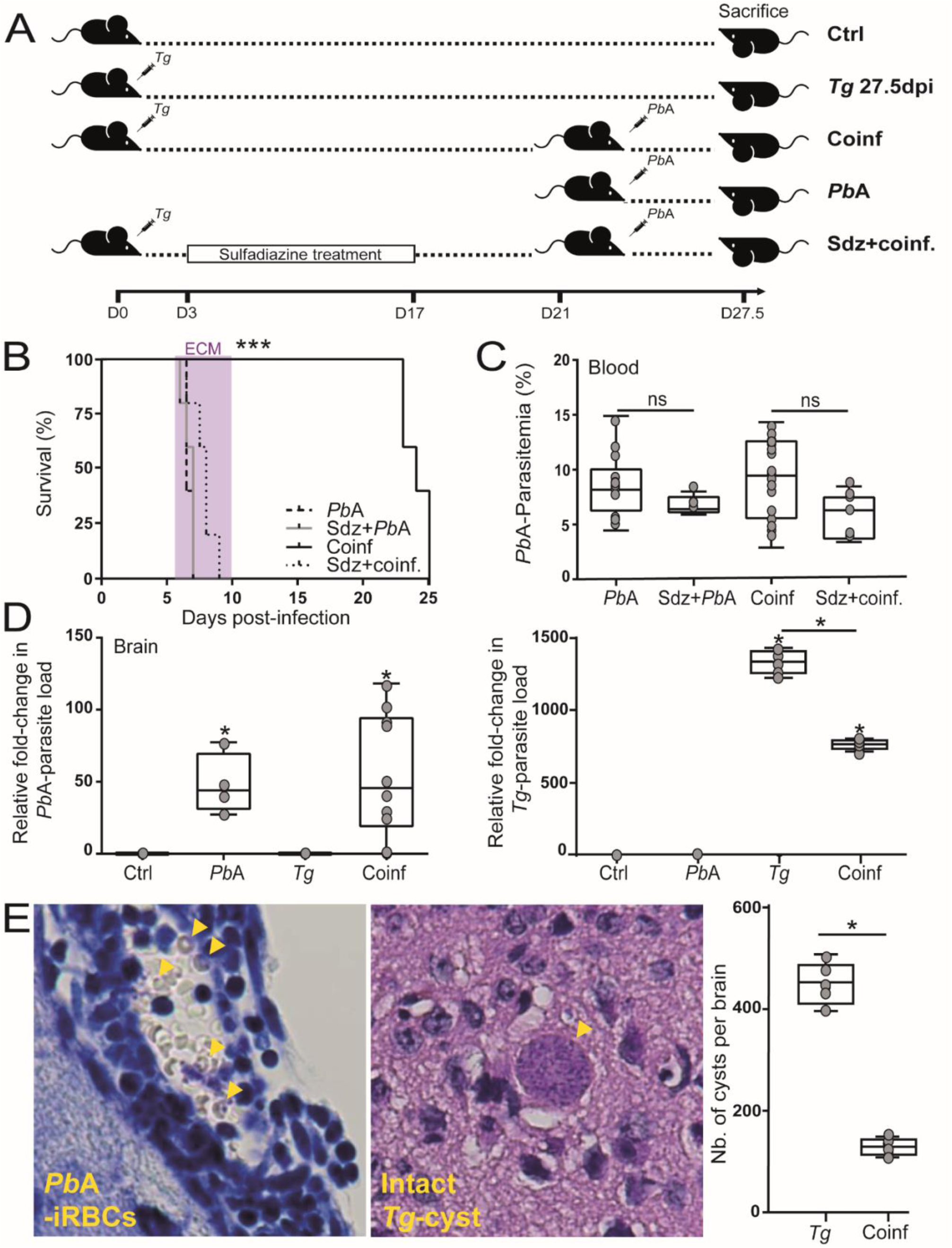
Model of *Tg*-*Pb*A coinfection C57BL/6 mice. (**A**) 6 weeks-old B6 mice were infected with 10^3^ *Tg* Type II Pru-GFP tachyzoites at day 0. At the day 21 (formation of cerebral cysts), mice were injected with 10^6^ blood stages of *Pb*A. Groups infected with *Tg* or *Pb*A alone and naive mice were used as controls in all experiments. *Tg* infected mice were also treated with SDZ (at 1g/l in drinking water) during 14 days after *Tg* infection (from day post-infection 3 to 17). (**B**) Survival rate of *Pb*A- or *Tg*-infected only and coinfected mice, treated or not with SDZ were followed daily. *Pb*A-mice group died during the ECM window, between 6 and 10 dpi (in violet). Statistically significant difference in survival curve, analyzed by the Log-Rank test, was observed in coinfected mice group without SDZ treatment compared to all other infected mice group. (**C**) Time course of *Pb*A-infection was monitored by assessing peripheral parasite in blood during onset of ECM around 6.5 dpi in all experimental groups. Parasitemia was assessed by microscopy of Giemsa-stained thin blood smears. (**D**) Parasite loads in brain during ECM and chronic *Tg* infection in all experimental groups. *Pb*A load in the whole brain was measured by *Pb18s* gene expression and *Tg* load by parasites genomes/µg brain DNA, using quantitative RT-PCR. (**E**) Histochemistry of brain sections showed presence of *Pb*A-iRBCs in microvessels and *Tg*-cyst (yellow arrows) in the cerebral parenchyma. Enumeration of cysts was done using microscopy by quantification of FITC-fluorescence labelling the wall of parasitic cysts. Values were expressed as fold-change compared to non-infected mice. All data were done on n = 4-8 samples from three to four independent experiments. Data were expressed as mean ± SEM and compared between groups, using a One-way ANOVA with Tukey’s multiple comparison test. Student’t test was used for two groups comparison. Ns = no significant; *P<0.05; ***P<0.001.

### Coinfection model

Five groups of mice were studied: (i) uninfected controls, (ii) *Pb*A-only, (iii) *Tg*-only, (iv) *Tg*–*Pb*A coinfected mice (infected with *Tg* for 21 days before *Pb*A challenge, as described by Settles *et al*. 2014), and (v) *Tg*–*Pb*A mice treated with sulfadiazine (SDZ) to inhibit cyst formation (Figure 1A) (33). All mice were sacrificed at 6.5 days post-*Pb*A infection, except for survival studies.

### RNA extraction and gene expression by real-time PCR

As described by Leleu *et al*., total RNA was isolated from whole brains using the NucleoSpin RNA kit (Macherey-Nagel; 750,955,250) and cDNA synthesis was performed with the SuperScript VILO kit (Invitrogen; 11,754,250) (12). From mix of cDNA with primers (Supplementary Table 1) (Eurogentec), RT-qPCR was carried out with SYBR Green Master Mix (Applied Biosystems; 4,385,612) on a QuantStudio 12K Flex system (ThermoFisher) to quantify cytokine/chemokine expression and parasitemia. Gene expression was normalized to *Hprt1* (Hypoxanthine phosphoribosyltransferase 1 gene) and analysed using the ΔΔCt method.

### Histology of brain sections

As described by Hellani *et al*., optimal cutting temperature compound (OCT) was used to embedded tissue of control and (co) infected mice (4). Coronal brain sections were cut serially at 5-µm thickness, spread onto Super-frost Plus slides (ThermoFisher, J1820AMNZ) and then allowed to dry vertically overnight at 37°C. To visualize parasites in cerebral parenchyma and to assess the degree of immune cell infiltration, vascularis and edema in meninges, brain sections were stained via haematoxylin and eosin (H&E). Slides were coverslipped using DPX mounting agent (Sigma-Aldrich). Slides were finally examined under light microscopy at 60X magnifications and images were taken using a digital camera mounted on the microscope.

### T_2_W images sequence Magnetic resonance imaging (MRI)

T-_2_ weighted (T_2_W) MRI sequences were performed on a Biospec 7.0T/20cm machine with a horizontal magnet (Bruker), as previously described by Dalko *et al* (9). At day 27.5, each group of animals were anesthetized with isoflurane and then placed in a dual-coil small animal restrainer containing a volume coil for transmission and a surface quadratic coil for reception of signals. Coil-to-coil electromagnetic interaction was actively decoupled. Respiration rates and waveforms were continuously monitored *via* a force transducer. A feedback-regulated circulating water pad was used to maintain the rectal temperature at 37±1°C. The acquisition of anatomic T_2_W images were done in axial and sagittal planes *via* the fast spin-echo TurboRARE (rapid acquisition relaxation-enhanced) pulse sequence, specifically configured (repetition time: 5000ms; echo time: 77ms; matrix size: 256×256 pixels; field of view: 20×20mm; number of excitations: 4). Acquisition time was 10min for each T_2_W sequence and choice of non-overlapping 0.5mm-thick slices was done to acquire axial and sagittal images. Apparent diffusion coefficient (ADC) was measured to estimated diffusion rates.

### Assessment of brain vascular permeability

BBB integrity was assessed 6.5 days post-*Pb*A infection using the Evans blue assay (Sigma-Aldrich; E2129), as described by Keswani *et al* (11). Brains were perfused, incubated in formamide, and dye extravasation quantified spectrophotometrically at 620 nm. Using a standard curve, Evans blue concentration was calculated and expressed per g of brain tissue. Visually, blue coloration of the tissue was a sign of brain vascular permeability.

### Cytokines and chemokines assay

Serum levels of IL-6, IL-10, IL-12, IL-33, TNF-α, CCL-2, and CXCL-10 were measured using a multiplex “MSD U-PLEX Mouse assay” (Meso Scale Discovery; K15069L-1). TGF-β levels were quantified by ELISA (R&D Systems; DY1679-05).

### Isolation of total brain cells

At day 27.5, brains were dissociated using a neural tissue dissociation kit (Miltenyi Biotec, 130-093-231), as described by Keswani *et al* (11). Then, Cells were purified by Percoll (Cytiva; 11748498) gradient centrifugation, erythrocytes lysed with ammonium-chloride-potassium (ACK) buffer, and viability assessed by trypan blue exclusion (Gibco; 15250061).

### Extra-/intra-cellular staining and flow cytometry

Cells were firstly treated with brefeldin A at 0.01 mg/mL (Sigma; B6542) for 4 h at 37°C. After 1X Phosphate-Buffered Saline (PBS) (Gibco; 10010023) washes, total brain cells were stained 30minutes at 4°C in 1X PBS (Gibco; 10010023)-10% FBS (Dutscher; S181B-500) with primary antibodies targeting extracellular and lineage markers (Supplementary Table 2). Cells were washes twice, then permeabilized and fixed using the CytoPerm-CytoFix kit (BD Biosciences; AB_2869008) 20 minutes at 4°C, and washed again. Intracellular staining was done by incubation with antibodies cocktail, in accordance to dilution stated in Supplementary Table 2, in 1X PBS (Gibco; 10010023) -10% FBS (Dutscher; S181B-500), overnight at 4°C (Supplementary Table 2). Finally, cells were ready after twice washes, or after a last step of streptavidin staining if needed, and resuspended in in 1X PBS (Gibco; 10010023) -10% FBS (Dutscher; S181B-500). Data of 1–5×10^5^ events were acquired using the LSR Fortessa (Becton Dickinson), after register compensations with monostaining. The different study populations were gating following their interest markers and the results were analysed on FlowJo software (version 10.9).

### Confocal microscopy of brain sections

Immunofluorescence staining was performed as described in Hellani *et al* (4). Briefly, coronal brain sections (5 μm, OCT-embedded; FisherScientific; 11912365), spread into Super-frost Plus slides (ThermoFisher; J1820AMNZ), were fixed in 4% paraformaldehyde (PFA; Electron Microscopy Sciences; 15,713 R7G5), permeabilized with 0.1% Triton X-100 (Sigma-Aldrich; T8787), and blocked (30 min in 1X PBS-5% FBS (Dutscher; S181B-500) prior to immunofluorescence staining. Primary antibodies targeted GFAP (glial fibrillary acidic protein; 1/400; ThermoFisher; 53-9892-80), IL-33 (1/200; R&D systems; AF3626), ST2 (1/200; Biolegend; 165302), and GATA3 (GATA-binding protein 3; 1/100; Santa Cruz Biotechnology; sc-268 AF594), followed by appropriate secondary antibodies (anti-mouse IgG-AF633 (1/200; Invitrogen; A21050); anti-mouse IgG-AF555 (1/200; Invitrogen; A31570)) and DAPI nuclear counterstain (Invitrogen; D1306). Images were acquired on a Zeiss LSM710 confocal microscope (ZEISS microscopy GmbH equipped with Airyscan super-resolution (0.325µm/pixel).

### Statistical analysis

Each experiment was performed at least twice with three to ten animals per group. GraphPad Prism (Version 9; GraphPad software, Inc.) was used for statistical analyses, with data expressed as a mean or median ± standard error of the mean (SEM). The two groups were compared using the unpaired Student’s t test and analysis of variance (ANOVA), followed by Tukey’s multiple comparison, were used to compare control and infected groups. Comparison of survival were done using the Log-Rank (Mantel Cox) test. Hierarchical clustering of the samples and cytokines/chemokines quantification were generated based on the Euclidean distance, using Perseus software (Max Planck Institute of Biochemestry) and expressed on heatmap after Y values transformation as Y=Log2(Y) with Prism. *P* values indicate statistical significance as *P<0.05, **P<0.01, ***P<0.001, and ****P<0.0001.

## Results

### 1. *Tg*-mediated protection against ECM requires the establishment of bradyzoite cysts in the brain without altering *Pb*A-iRBC sequestration in microvessels

To investigate the mechanisms underlying the modulation of *Pb*A-induced neuropathogenesis during coinfection with *T. gondii* (*Tg*), we adapted the model previously described by Settles *et al*. (33). C57BL/6 mice were first infected with type II *Tg* (Pru strain) tachyzoites for 21 days, leading to the establishment of bradyzoite cysts in the brain, as confirmed by RT-qPCR (data not shown). Mice were then coinfected with 10^6^ blood stage of *Pb*A clone 1.4 known to induced ECM (Figure 1A). Mice infected with *Tg* or *Pb*A alone and naive mice were used as controls in all experiments. An additional group of *Tg*-infected mice received sulfadiazine (SDZ) treatment at day 3 post-infection to eliminate tachyzoites prior to brain invasion, while preserving the early systemic pro-inflammatory response (Figure 1A). As expected, *Pb*A-infected mice, whether or not pre-treated with SDZ, succumbed to ECM by day 6.5 (pink window in figure 1B). In contrast, chronically *Tg*-infected mice were fully protected from ECM upon *Pb*A challenge, although they later succumbed to hyperparasitemia at 23– 24 dpi (Figure 1B). Importantly, SDZ treatment abrogated the protective effect of *Tg* infection, indicating that the establishment of cerebral *Tg* cysts was required for ECM resistance. Coinfection did not alter *Pb*A parasitemia at 6.5 dpi (Figure 1C), and comparable *Pb18s* (18S ribosomal RNA gene of *Plasmodium berghei*) transcript levels were detected in *Pb*A-only and *Tg*–*Pb*A groups (Figure 1D), demonstrating that parasite fitness was unaffected. Conversely, *Tg* mRNA levels were reduced in the brains of *Tg*–*Pb*A coinfected mice compared to *Tg*-only mice (Figure 1D). Finally, histochemical analysis confirmed the presence of *Pb*A-iRBCs within cerebral microvessels and *Tg* cysts in the parenchyma. Cyst quantification, performed by FITC fluorescence microscopy of cyst walls, revealed that *Pb*A sequestration persisted in the brains of coinfected mice, indicating that ECM protection was not linked to reduced iRBC accumulation. Nevertheless, *Tg*–*Pb*A coinfected mice displayed fewer cerebral cysts at 27 dpi compared to *Tg*-only mice, in which cysts remained detectable without associated immune infiltration (Figure 1E).

### 2. BBB damages and cerebral edema are prevented in *Tg*-*Pb*A coinfected mice

We assessed physiological and morphological changes in the CNS of *Pb*A-infected mice during ECM, as well as in *Tg*-only and *Tg*–*Pb*A coinfected mice, using MRI. T_2_-weighted (T_2_W) images revealed pronounced alterations in the corpus callosum of *Pb*A-infected mice (yellow boxe in control images), consistent with edema and hyperintense lesions indicative of structural damage typically associated with ECM (Figure 2A). Volumetric analysis confirmed a significant increase in corpus callosum volume in *Pb*A-infected mice compared to non-infected controls and *Tg*-infected groups (Figure 2B). In contrast, both *Tg* groups exhibited preservation of corpus callosum integrity, with volumes comparable to controls (Figure 2A–B). Analysis of ventricular morphology showed distinct patterns in the third ventricle (yellow arrows): *Tg*-only and *Tg*–*Pb*A mice exhibited increased visibility primarily in the upper portion, whereas ECM^+^ mice displayed enlargement predominantly in the lower portion (Figure 2A), reflecting differential structural remodelling and highlighting distinct neuropathological signatures between *Tg*- and *Pb*A-driven cerebral alterations. In addition, MRI assessment of the olfactory bulbs (yellow ellipse in figure 2A) demonstrated marked volumetric expansion across all infected groups compared to uninfected controls (Figure 2B). Notably, co- and *Tg*-only infected mice brain presented approximately two-fold larger olfactory bulbs than ECM^+^ group. No significant difference was observed between *Tg*-only and *Tg*–*Pb*A groups, indicating that *Tg* infection, independently of *Pb*A coinfection, drives this pronounced olfactory bulb hypertrophy. Diffusion-weighted imaging revealed reduced ADC values in cortical regions of *Tg*-only and coinfected mice compared to *Pb*A group, suggesting lower brain inflammation than observed during ECM (Figure 2C). Consistently, BBB) permeability assessed by Evans blue dye infiltration was increased in ECM^+^ mice but remained unaltered in *Tg*–*Pb*A coinfected mice, indicating preserved BBB integrity during coinfection (Figure 2D). To finish, histological analysis confirmed the MRI observations. *Pb*A-infected mice displayed pronounced parenchymal immune cell infiltration and vasculitis, whereas these features were markedly reduced in *Tg*–*Pb*A coinfected mice. All infected groups exhibited meningeal immune infiltration and extracellular edema, with meningeal thickening present in both *Pb*A- and *Tg*-infected animals compared to controls. Notably, parenchymal inflammation and vasculitis were attenuated in coinfected mice relative to *Tg*-only mice (Figure 2E). Together, these results demonstrate that *Pb*A infection primarily induces ECM-associated white matter swelling, whereas *Tg* infection leads to olfactory bulb hypertrophy and reduced neuroinflammation. Protection against ECM during coinfection with *Tg* is thus linked not to impaired *Pb*A-iRBC sequestration but to modulation of neuroinflammatory responses.

**Figure 2.**
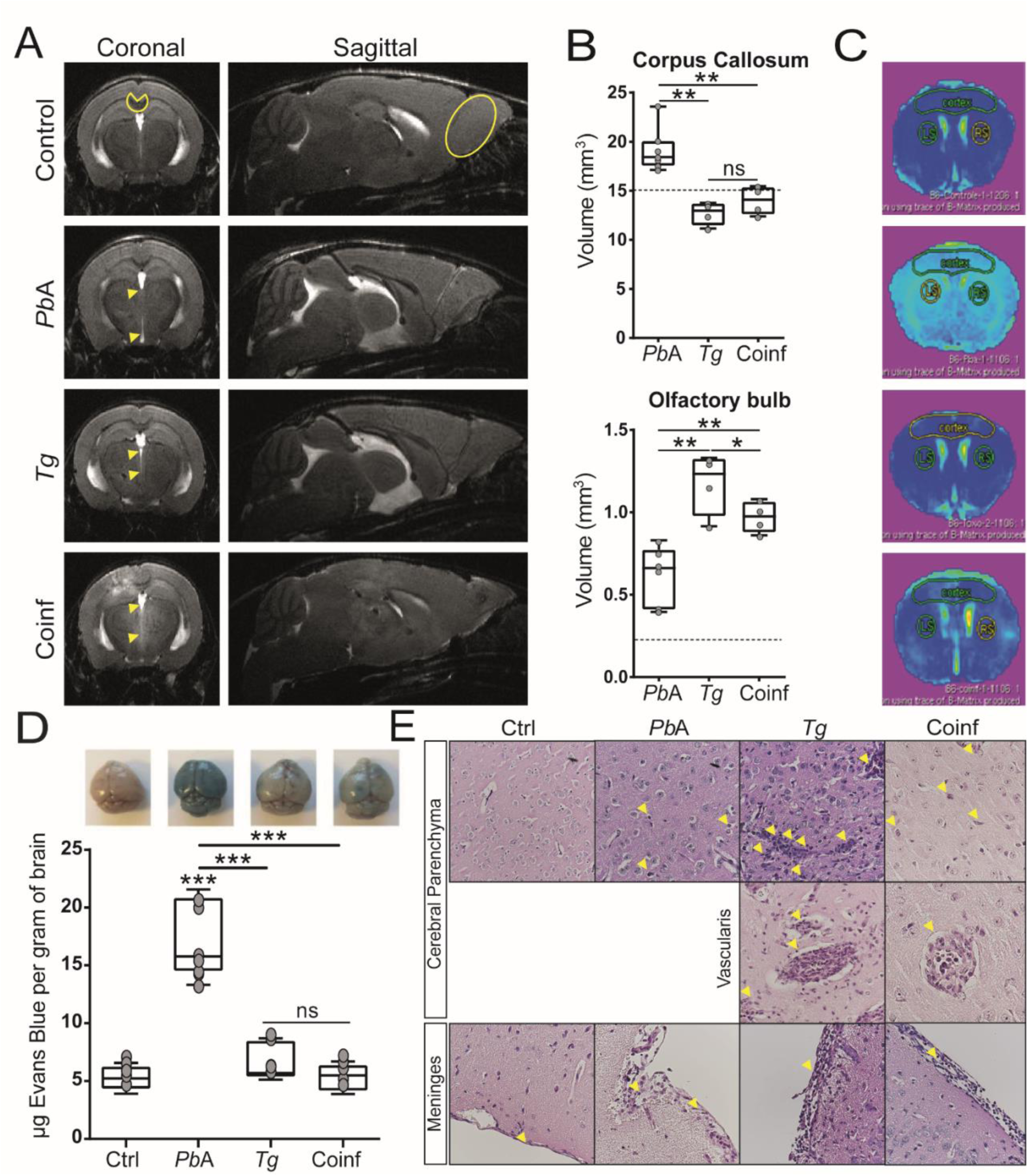
Coinfected mice show reduced brain edema and preserved BBB integrity. (**A, B**) Coronal and sagittal T_2_-weighted MRI images of control, ECM^+^, *Tg*-only, and *Tg*–*Pb*A mice. Coinfected mice display less disruption in the corpus callosum and olfactory bulbs compared to ECM^+^ mice. Yellow boxe indicate hyperintense corpus callosum regions, ellipse mark olfactory bulbs, and yellow arrows show third ventricle enlargement. Student’s t-test, P<0.01. (**C**) ADC maps reveal reduced cortical inflammation in coinfected mice versus ECM^+^. (**D**) Evans blue assay shows BBB leakage only in *Pb*A-infected mice. Data: mean ± SEM; one-way ANOVA with Tukey’s test. (**E**) Histology of brain sections. Coinfected mice exhibit reduced parenchymal immune infiltration and vasculitis (yellow arrows) compared to *Tg*-only mice. Meningeal thickening is present in all infected groups. ns = not significant; ***P < 0.001.

### 3. ECM protection involves TGF-β and IL-33–driven control of neuroinflammation

To investigate the molecular mechanisms underlying *Tg*-mediated protection against ECM, we quantified cytokines and chemokines in the brain of control, *Pb*A-, *Tg*-, and *Tg*–*Pb*A-infected mice by RT-qPCR (Figure 3A). Gene expression analysis revealed significant differences across groups, forming three distinct clusters. Both *Pb*A and *Tg* infection induced upregulation of pro-inflammatory mediators such as *Cxcl-10*, *Ccl-2*, *Il-6*, and *Tnf-α*, compared to controls, whereas *Tg-* and co-infection were uniquely associated with enhanced expression of the immunoregulatory genes *Il-10* and *Tgf-β*, consistent with attenuated neuroinflammation compared to the severe response observed during ECM. Examination of the IL-33 pathway further revealed that *Tg* infected groups exhibited strong upregulation of *Il3-3a* and its receptor *St2*, but not of inducible *Il-33b*, in contrast to ECM^+^ mice where expression remained moderate (Figure 3A). Notably, while *Cxcl-10* expression was increased in all infected groups, it was only modestly higher in brain of *Tg* groups (∼1.2-fold), whereas *Tg* infection specifically drove robust induction of *Tgf-β*, *Il-33a*, and *St2* (∼5-, ∼3-, and ∼6-fold, respectively) compared to *Pb*A mice brain (Figure 3B). Serum cytokine profiles further supported these findings, distinguishing two patterns: (i) low or moderate systemic inflammation in controls and chronic *Tg* infection, and (ii) high systemic inflammation in *Pb*A and *Tg*–*Pb*A groups (Figure 3C–D). Unlike in the brain, secreted IL-33 was elevated in the sera of all infected mice (Figure 3D). Together, these results indicate that ECM is driven by an exacerbated CXCL-10-mediated pro-inflammatory response, whereas *Tg* infection promotes TGF-β and IL-33/ST2 signalling, which may mitigate ECM-associated neuroinflammation during coinfection.

**Figure 3.**
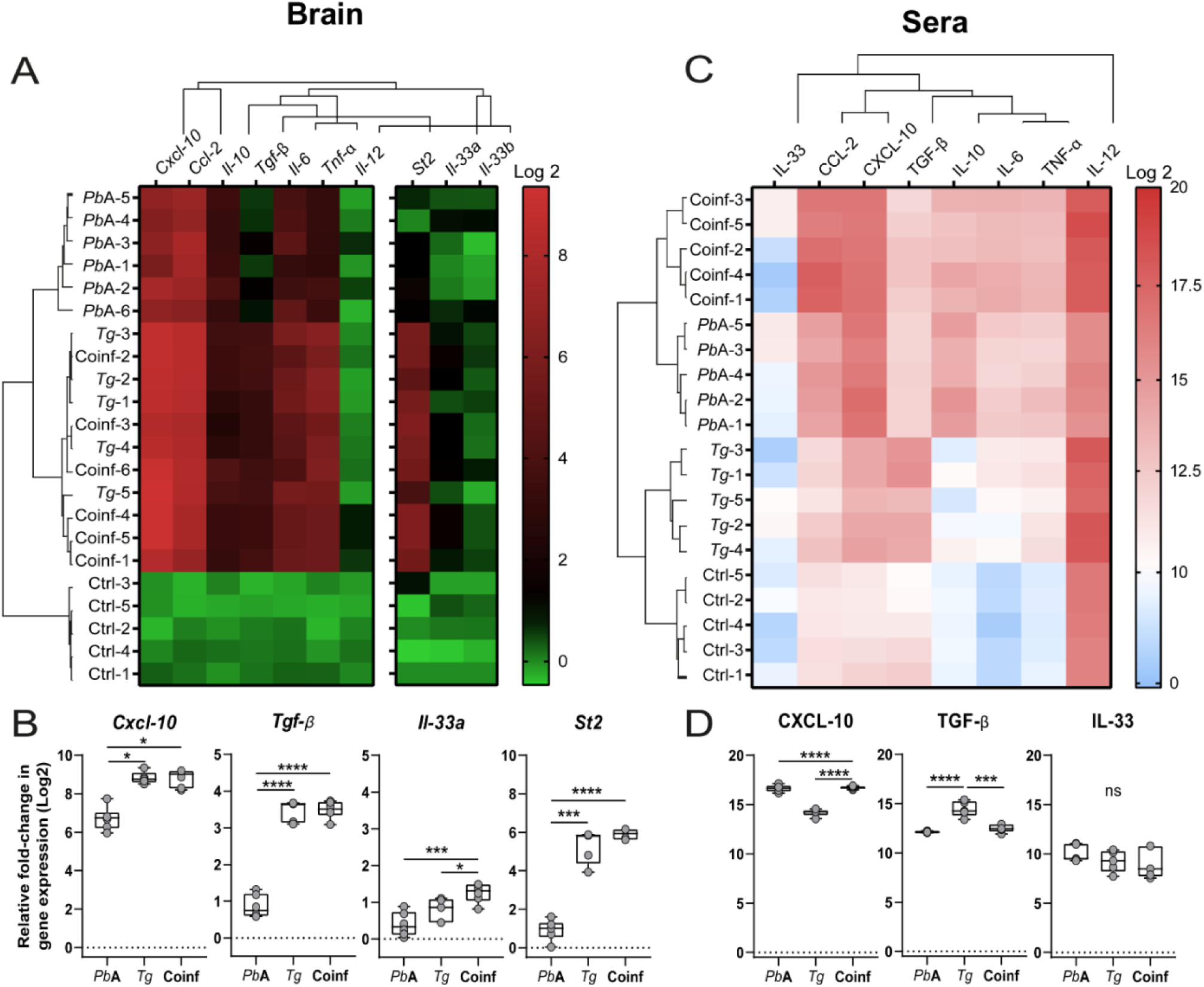
Protection against ECM associated with a neuroinflammation controlled by TGF-β and IL-33/ST2 pathway. Clustering of gene relative expressions (A,B) and systemic levels (C,D) of a panel of pro- and anti-inflammatory cytokines and chemokines, involved in the ECM and the chronic *Tg*. Data were measured, by RT-qPCR, in the brain and by multiplex ELISA, in the sera of mice infected by *Pb*A and/or *Tg*, compared to controls. Values (n=5-6) were transformed and represented using Y=Log2(Y). Box show all points, between the minimum and the maximum values, and the median ± SEM. Statistical significance was determined using a One-way ANOVA with Tukey’s multiple comparison test. P values indicate statistical significance as at ns= no significant; *P<0.05; ***P<0.001; ****P<0.0001.

### 4. Coinfection alters astrocyte and microglia inflammatory responses

We next analysed brain immune cell composition across all groups by flow cytometry. Total cell counts were significantly increased during both *Tg* and *Pb*A infection, with the strongest effect observed in *Tg*-infected mice. In contrast, coinfected mice showed reduced numbers of total brain cells compared to *Tg* chronic infection (Figure 4A). We then quantified astrocytes (GLAST^+^ glutamate aspartate transporter positive; GFAP^+^ positive); and microglia (CD11b^+^ CD45^low^ cells). *Tg* infection markedly induced astrogliosis increasing the number of GLAST⁺ astrocytes, while *Pb*A had little effect (Figure 4B). The proportion of GFAP⁺ astrocytes increased in all infected groups relative to controls. *Tg* infection induced high Gmean activation in ∼8% of astrocytes, while ECM was associated with a much stronger activation (∼30%). Interestingly, *Pb*A infection in chronically *Tg*-infected mice did not exacerbate astrocyte activation, being reduced (∼5%) compared to *Tg* chronic infection (Figure 4C). This trend was confirmed by quantification of Major Histocompatibility Complex (MHC)-I⁺ astrocytes, although MHC-I expression levels (fold Gmean) remained similar across ECM, *Tg*, and coinfected groups (Figure 4D). We further examined the inflammatory phenotype of reactive astrocytes by assessing CXCL-10 (A1 pro-inflammatory) and TGF-β (A2 anti-inflammatory) production in GLAST⁺GFAP⁺ cells. In ECM, astrocytes predominantly produced CXCL-10, whereas *Tg* infection favoured TGF-β expression (Figure 4E-F). Remarkably, coinfected mice displayed concomitant production of CXCL-10 (∼10%) and TGF-β (∼15%) with moderate Gmean, indicating the emergence of an intermediate A1/A2 phenotype (Figure 4E-F).

**Figure 4.**
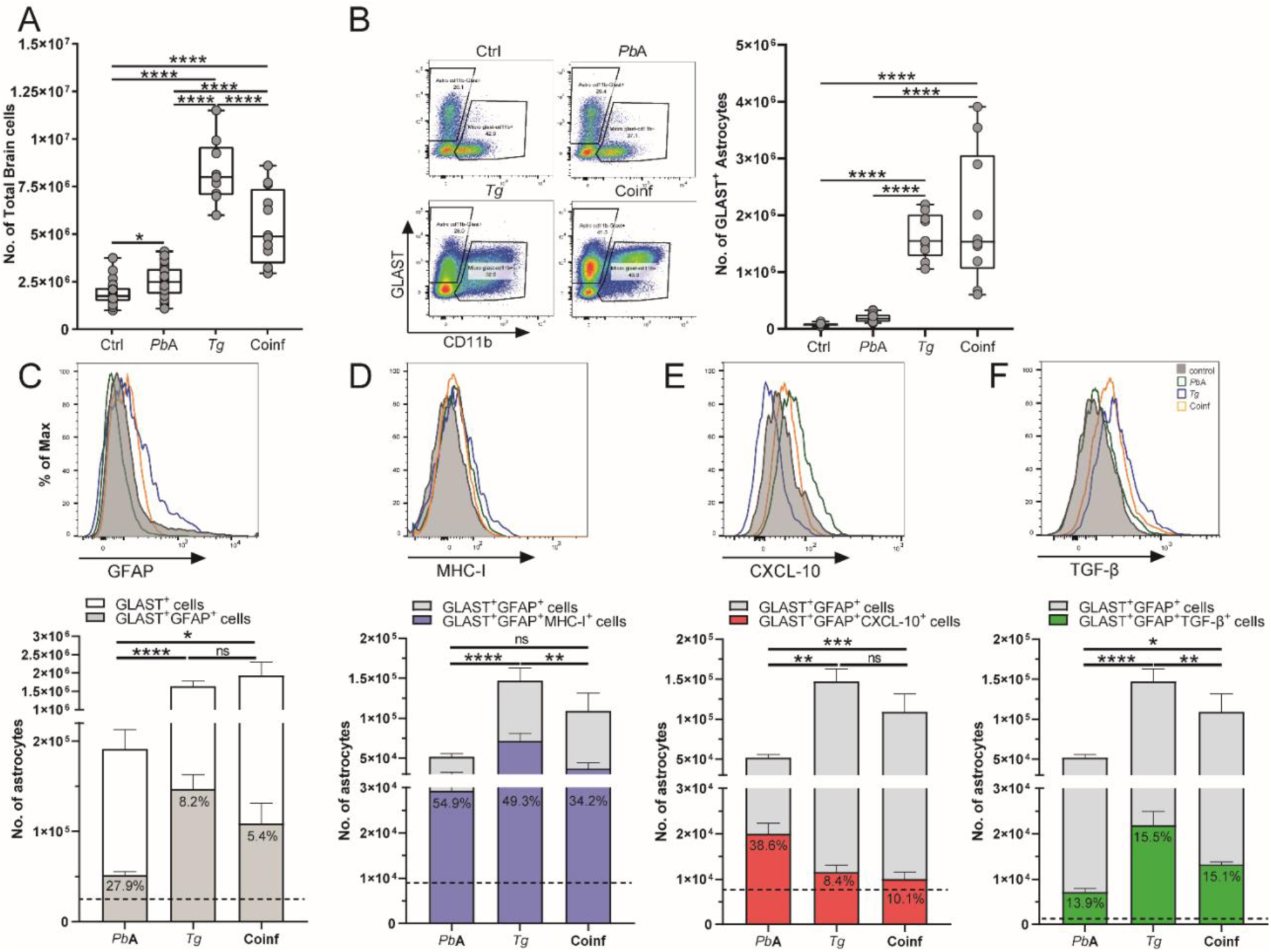
Changed activation and inflammatory phenotype of astrocytes during *Tg*–*Pb*A coinfection. Flow cytometry analysis of total brain cells (**A**) and GLAST^+^ astrocytes (**B**) showing increased numbers during *Tg* infection compared to uninfected and *Pb*A-infected controls. (**C**) Quantification of GFAP⁺GLAST⁺ astrocytes revealed reduced activation in *Tg* and coinfected groups relative to ECM⁺ mice. Frequency (% of Max) plots confirmed that only a small fraction of astrocytes expressed GFAP in coinfected brains, contrasting with *Tg* infection. Within this population, MHC-I expression (D) and production of CXCL-10 (E) and TGF-β (F) were assessed. Coinfected astrocytes displayed lower MHC-I expression than *Tg*- and ECM⁺ mice *Tg* infection, produced more CXCL-10 but less TGF-β than A2 in chronic *Tg*. Data are presented as median ± SEM (n = 10, five independent experiments). One-way ANOVA with Tukey’s multiple comparisons test was used; ns, not significant; *P < 0.05; **P < 0.01; ***P < 0.001; ****P < 0.0001.

We further compared the phenotypes of microglia in *Tg*-*Pb*A coinfected with *Tg-* and ECM^+^ mice (Figure 5). Similar to astrocytes, the number of CD11b^+^CD45^low^ microglia was increased in all infected groups relative to controls, with the strongest microgliosis observed in *Tg*-infected mice compared to ECM^+^ animals (Figure 5A). However, this expansion was less pronounced in coinfected mice (Figure 5B). We then assessed microglial activation states by evaluating pro-inflammatory (M1) markers -CD86, CD16/32, and MHC class I/II- and the anti-inflammatory (M2) marker CD206 (35,36). During ECM, microglia predominantly exhibited an M1 profile (CD86^hi^ CD16/32^hi^ MHC-II^low^) with moderate CD206 expression (Figures 5B–C). By contrast, chronic *Tg* infection induced a distinct profile with lower expression of CD86 (∼1.2-fold), CD16/32 (∼1.3-fold), and CD206 (∼2.1-fold) compared to *Pb*A, but with slightly higher MHC-II expression (∼1.3-fold). Notably, CD206 was strongly expressed in *Tg*-infected mice (Figure 5C). Coinfected mice displayed an M1-like profile similar to ECM, but with markedly elevated MHC-II expression (Figure 5C). Focusing on CD86⁺ microglia, we found that coinfection was associated with reduced MHC-I expression and enhanced CXCL-10 production, as during ECM, compared to chronic *Tg* (Figures 5D–E). Conversely, CD206⁺ microglia exhibited comparable MHC-I expression across *Pb*A, *Tg*, and coinfected groups, but *Pb*A superinfection unexpectedly triggered high TGF-β production by this population (Figure 5F–G).

**Figure 5.**
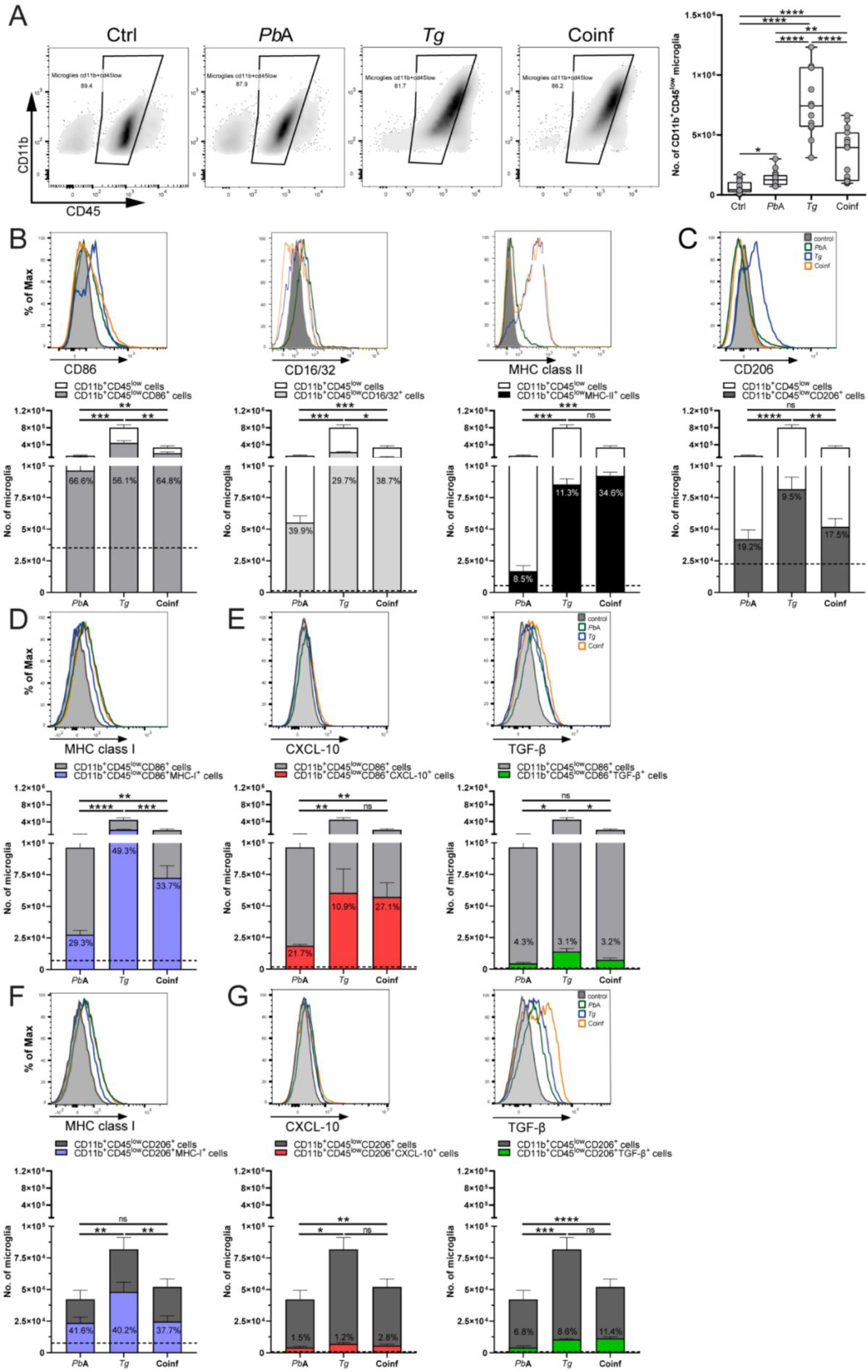
Intermediate microglial inflammatory phenotype during *Tg*–*Pb*A coinfection. Flow cytometry quantification of CD11b⁺CD45^low microglia in *Tg*-, *Pb*A-, and *Tg*–*Pb*A-infected mice compared to uninfected controls (**A**). Quantification by flow cytometry of CD11b^+^CD45^low^ microglia in brain of *Tg-* or *Pb*A- or *Tg*-*Pb*A coinfected mice compared to uninfected. Frequency and total number of CD11b^+^CD45^low^ activated microglia expressing M1 markers (**B**) as CD86, CD16/32 and MHC class II, and M2 marker (**C**), CD206, were represented in the total number of microglia, in all mice groups. Majority of microglia in brain of coinfected mice highly expressed all of them. Then, expression of MHC class I and production of CXCL-10 and TGF-β (frequency and total number) were compared in CD86^+^M1 phenotype (**D,E**) and CD206^+^M2 phenotype (**F,G**) cells. Data were represented as median ± SEM (n = 10, from five independent experiments) and compared between groups. Statistical significance was determined using a One-way ANOVA with Tuckey’s multiple comparison test. P values indicate statistical significance as at ns= no significant; *P<0.05; **P<0.01; ***P<0.001; ****P<0.0001. Data are presented as median ± SEM (n = 10, five independent experiments). Statistical significance was determined by one-way ANOVA with Tukey’s multiple comparisons; ns, not significant; *P < 0.05; **P < 0.01; ***P < 0.001; ****P < 0.0001.

Together, our results demonstrate that ECM induces strong pro-inflammatory activation of astrocytes and microglia (A1/M1), while chronic *Tg* infection promotes an anti-inflammatory profile (A2/M2). During *Tg*–*Pb*A coinfection, both cell types adopt an intermediate A1/A2–M1/M2 phenotype, producing CXCL-10 and TGF-β. This balanced gliosis likely limits neuroinflammation and underlies protection against ECM.

### 5. Monocyte recruitment in the brain during *Tg*–*Pb*A coinfection

We next analysed brain monocyte subpopulations across all experimental groups (Figure 6). Total CD11b^+^CD45^hi^ monocytes increased in all infected mice, with the highest numbers observed in *Tg*– *Pb*A coinfected animals (Figure 6A). Subdivision by Ly6C (Lymphocyte antigen 6) expression (Figure 6B) revealed expansion of Ly6C^hi^ classical resident, Ly6C^int^ intermediate, and Ly6C^low^ non-classical patrolling monocytes, particularly during coinfection (Figures 6C-E). Next, we especially examined non classical “patrolling” (Ly6C^low^Ly6G^-^CCR2(C-C Chemokine Receptor type 2)^-^ CX3CR1(C-X3-C Chemokine Receptor 1)^hi^CD62L^-^), resident intermediate (Ly6C^int^Ly6G^-^ CCR2^int^CX3CR1^int^CD62L^+^) and inflammatory (Ly6C^hi^Ly6G^-^CCR2^hi^CX3CR1^-^CD62L^+^) monocytes. Our data showed that a huge number of patrolling and resident intermediate and inflammatory monocytes are recruited in the brain during the concomitant infections with *Pb*A and *Tg* (Figures 6F-G). Patrolling Ly6C^low^ monocytes were largely absent in ECM^+^ mice but increased in *Tg*-infected and coinfected mice, indicating recruitment driven by chronic *Tg* rather than *Pb*A (Figure 6F). This suggest that patrolling monocytes were recruited when the *Tg* infection starts and their numbers were not influenced by the *Pb*A secondary infection. It is known that Ly6C^+^ monocytes are necessary for the early control of *Tg* replication (37). Intermediate Ly6C^int^ monocytes were abundant in *Tg* and coinfected brains, with coinfection having no significant effect on their numbers (Figure 6G). This suggests that infection with *Pb*A does not affect the recruitment of the Ly6G^int^ intermediate monocytes, suggesting to be induced by the chronic *Tg*. Regarding the Ly6^hi^ inflammatory monocytes, we find this population elevated in ECM^+^ and chronic *Tg* mice, and further amplified in coinfected mice, suggesting enhanced recruitment during chronic *Tg* infection reinforced by *Pb*A (Figure 6H). However, it was noted that the mice infected at 6.5 days with *Tg* (mice with acute toxoplasmosis), did not show a significant difference in quantity of inflammatory monocytes compared to control mice (data not shown). These results indicate that *Tg*–*Pb*A coinfection promotes selective recruitment of monocyte subsets and a controlled inflammatory response, likely contributing to neuroprotection during ECM.

**Figure 6.**
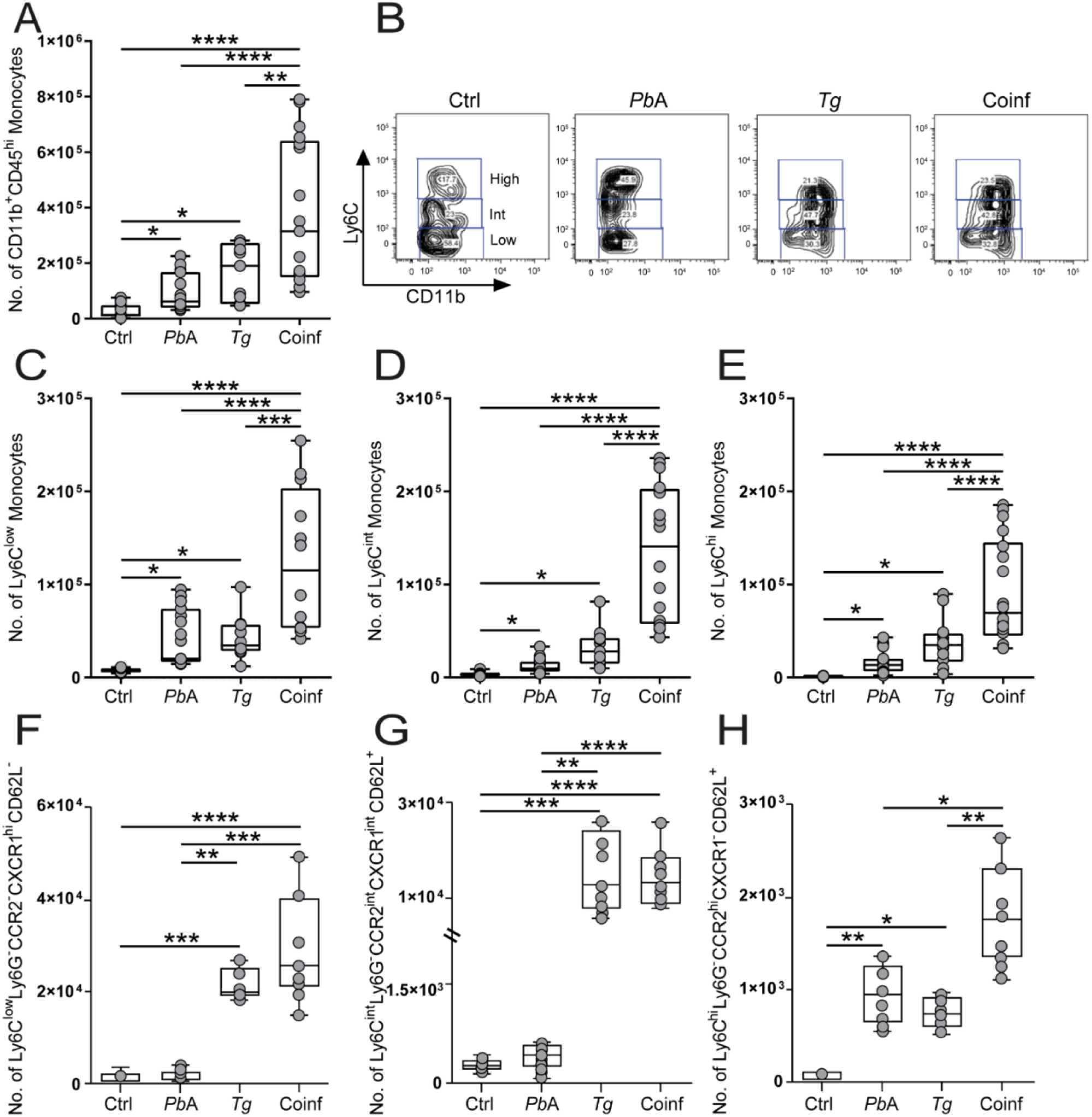
High monocytes infiltration in brain during the coinfection. (**A**) Increase of CD11b^+^CD45^hi^ monocytes in brain of *Tg* and/or *Pb*A infected compared to uninfected mice, quantified by flow cytometry. (**B**) Gating and total number of classical Ly6^hi^ (**C**) or not Ly6C^int^ (**D**)/Ly6C^low^ (**E**) monocytes, resident or infiltering the brain of mice with either protozoan, compared to control. Number of (**F**) patrolling (Ly6C^low^Ly6G^-^CCR2^-^CX3CR1^hi^CD62L^-^), (**G**) intermediate (Ly6C^int^Ly6G^-^CCR2^int^CX3CR1^int^CD62L^+^) and (**H**) inflammatory (Ly6C^hi^Ly6G^-^CCR2^hi^CX3CR1^-^CD62L^+^) monocytes more higher during coinfection by contrast to the others groups. Data were represented as median ± SEM (n=10, from five independent experiments) and compared between groups. Statistical significance was determined using a One-way ANOVA with Tuckey’s multiple comparison test. P values indicate statistical significance as at *P<0.05; **P<0.01; ***P<0.001; ****P<0.0001.

### 6. Reactive astrocytes drive ILC2s recruitment in the brain of *Tg*-*Pb*A coinfected mice, by the IL-33/ST2 pathway

During chronic *Tg* infection, astrocytes promote a protective immune response via the IL-33/ST2 pathway, driving ILC2 infiltration into the brain (31,32). We assessed the contribution of astrocyte-derived IL-33 to ECM protection during *Tg*–*Pb*A coinfection using IL-33 KO mice. Survival analysis showed no difference between B6 and IL-33 KO mice: both succumbed to ECM between 6–9 dpi when infected with *Pb*A, whereas *Tg*–*Pb*A coinfected mice died later from hyperparasitemia (Figure 7A). Flow cytometry revealed that GFAP⁺ astrocytes produced IL-33 during *Tg* infection and coinfection, at higher levels in *Tg* mice, whereas *Pb*A reduced IL-33 production during coinfection (Figure 7B). Notably, IL-33⁺ astrocytes co-expressed ST2 in *Tg* and coinfected mice, accounting for nearly one-third of activated astrocytes in coinfected brains (Figure 7C), which was confirmed by confocal microscopy (Figure 7D). We next analysed ILC2 infiltration, defined as Lineage^-^IL-7^+^ROR(Retinoic acid receptor-related orphan receptor)γt^-^GATA3^+^ cells (Supplementary figure 1). As in chronic *Tg* infection, ILC2s accumulated in the brains of *Tg*–*Pb*A coinfected mice (∼2×10⁵ cells), whereas IL-33 KO mice displayed a ∼10-fold reduction (Figure 7E). Confocal microscopy confirmed the presence of GATA3⁺ST2⁺ ILC2s in the brains of coinfected B6 mice (Figure 7F). Finally, we examined cytokine production by infiltrating ILC2s. In *Tg* and coinfected brains, ILC2s expressed IL-5 and IL-13, with IL-13⁺ cells being more abundant than IL-5⁺ cells during coinfection (Figure 8). These findings suggest that astrocyte-derived IL-33 recruits ILC2s through the IL-33/ST2 axis, and that ILC2s -particularly IL-13 producers-may contribute to protection against ECM in the context of *Tg* infection.

**Figure 7.**
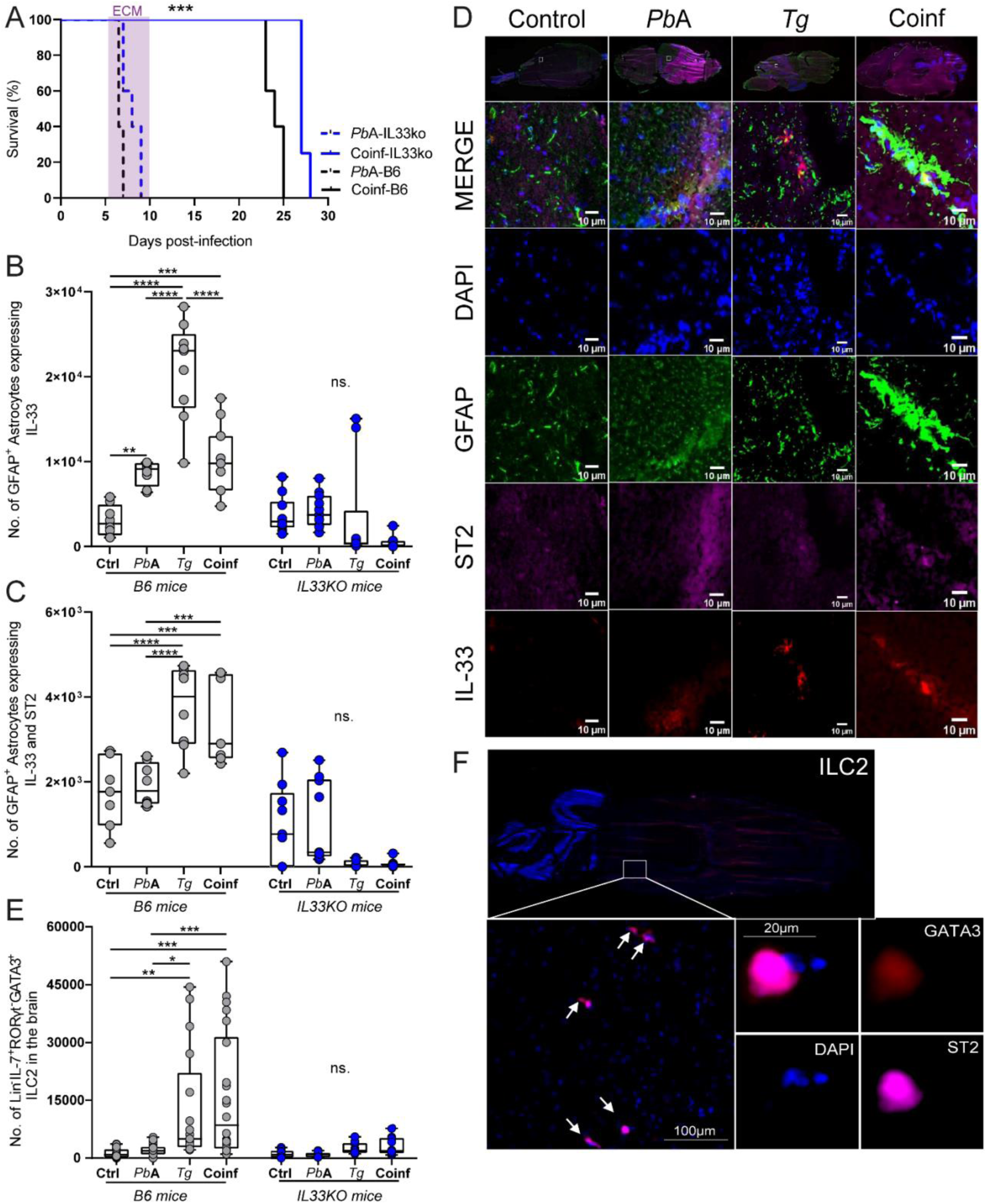
Astrocytes produced IL-33 which drives the ILC2s infiltration in the brain of *Tg*-*Pb*A coinfected mice. (**A**) Survival of IL-33 KO vs B6 mice, infected by *Pb*A or coinfected with *Tg*. Flow cytometry analysis of the number of GFAP^+^ activated astrocytes producing IL-33 (**B**) and ST2 (**C**) from brain of *Tg* or/and *Pb*A infected or not B6 and IL-33 KO mice. (**D**) IL-33 (red) and ST2 (pink) labels in GFAP^+^ astrocytes (green and DAPI in blue) in brain sections of B6 mice. (**E**) Number of ILC2s infiltrated in mice brain during *Tg* and coinfection with *Pb*A. (**F**) Observation of ILC2s co-expressing GATA3 and ST2 in brain section from *Tg*-*Pb*A coinfected B6 mice. Data were represented as median ± SEM (n = 3-10, from five independent experiments) and compared between groups. Statistical significance was determined using a One-way ANOVA with Tuckey’s multiple comparison test. P values indicate statistical significance as at ns= no significant; *P<0.05; **P<0.01; ***P<0.001; ****P<0.0001.

**Figure 8.**
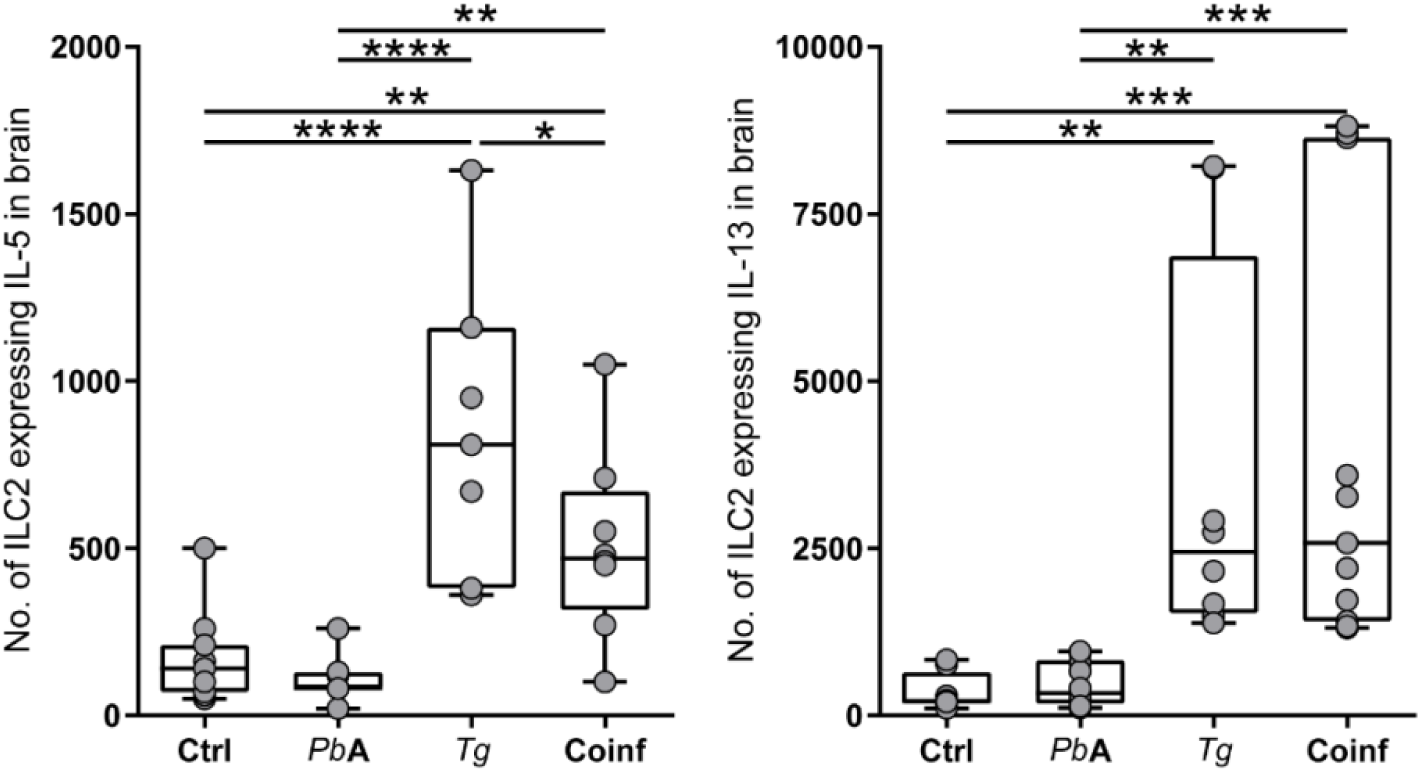
Expression of IL-5 and IL-13 by ILC2s infiltered in the brain of *Tg*-*Pb*A coinfected mice. ILC2s producing IL-5 and IL-13 were analyzed in cerebral cells isolated from the brain of control or infected mice by *Tg* and/or *Pb*A. By flow cytometry, these cells were then gated and quantified with the FlowJo software. High IL-13 and moderate IL-5 amounts were produced by ILC2s in coinfected groups. Data were represented as median ± SEM (n = 3-10, from five independent experiments) and compared between groups. A One-way ANOVA with Tukey’s multiple comparison test was used for statistics analysis and P values indicate statistical significance as at ns= no significant; *P<0.05; **P<0.01; ***P<0.001; ****P<0.0001.

## Discussion

Our comprehensive investigation of the coinfection dynamics between *Toxoplasma gondii* (*Tg*) and *Plasmodium berghei* ANKA (*Pb*A) provides critical mechanistic insights into the neuroprotective phenomena observed in experimental cerebral malaria (ECM). Employing the B6 experimental model, we demonstrate that the pathophysiological mechanisms underlying chronic *Tg* infection— characterized by the establishment of dormant bradyzoite-containing tissue cysts within neural parenchyma—exhibit remarkable convergence with the neuroinflammatory cascades implicated in ECM pathogenesis (25,29,38). This convergence provides the foundational basis for understanding how pre-existing chronic infection can fundamentally alter subsequent pathogenic challenges.

Our findings corroborate established paradigms of disease tolerance and immune regulatory networks operative in coinfection scenarios, demonstrating how chronic *Tg* colonization fundamentally reprograms the neuroinflammatory milieu within the CNS (39,40). This study elucidates the complex parasitic interactions that fundamentally alter host neuroimmune responses, ultimately determining divergent pathological trajectories and survival outcomes in polymicrobial infections. The observed neuroprotective phenotype was contingent upon the simultaneous cerebral presence of both parasitic species, establishing the critical requirement for active coinfection rather than sequential exposure. While *Tg* established characteristic tissue cysts and *Pb*A-infected erythrocytes sequestered within cerebral microvascular beds, coinfection paradoxically attenuated neuroinflammatory responses, as demonstrated through MRI. This spatial and temporal co-localization requirement indicates that neuroprotection emerges from direct parasitic interactions rather than systemic immune modulation alone. This contrasts markedly with the severe neuroinflammation and associated mortality observed in *Pb*A monoinfections, establishing the specificity of the protective mechanism to the coinfection state rather than general immune suppression.

Building upon the prerequisite spatial requirements, we identified specific molecular mechanisms underlying neuroprotection. Surviving coinfected mice exhibited a distinctive cytokine profile characterized by downregulated pro-inflammatory mediators (CXCL-10, CCL-2, IL-6, TNF-α) concurrent with upregulated anti-inflammatory cytokines (IL-10, TGF-β). This balanced immunoregulatory state prevents the catastrophic neuroinflammation associated with ECM pathogenesis while maintaining sufficient immune competence for pathogen control. The chemokine CXCL-10, typically hyperexpressed during *Pb*A infection and instrumental in orchestrating pathogenic CD8^+^ T cell neuroinvasion and ECM development demonstrated significantly moderated expression in coinfected animals (9,12,41). This attenuation represents a critical mechanistic component, as CXCL-10-mediated T cell recruitment constitutes a primary pathogenic pathway in ECM. The concurrent upregulation of TGF-β, a pleiotropic cytokine critical for tissue repair and fibrosis resolution, established a neuroprotective microenvironment that actively mitigates inflammation-induced neural damage (30). These observations align with emerging evidence demonstrating the critical roles of IL-10 and TGF-β in mitigating neuroinflammatory cascades and preserving neural tissue integrity, providing mechanistic validation for our observed protective phenotype (30,42–44).

The cytokine network remodeling described above is both mediated by and reflected in fundamental changes in glial cell activation states. Astrocytes, the predominant glial population within the CNS, demonstrate remarkable phenotypic plasticity in response to pathogenic stimuli, transitioning between neuroprotective (A2) and neurotoxic (A1) activation states (45). This cellular plasticity provides the mechanistic basis for the observed cytokine profile changes. *Pb*A monoinfection predominantly induces a neurotoxic A1 phenotype characterized by elevated CXCL-10 production, directly correlating with increased disease severity and mortality. This establishes the cellular source of the pathogenic inflammatory signals identified in our cytokine analysis. Conversely, *Tg* infection promotes a neuroprotective A2 phenotype distinguished by enhanced TGF-β secretion, providing the cellular mechanism for anti-inflammatory cytokine production. *Pb*A monoinfection predominantly induces a neurotoxic A1 phenotype characterized by elevated CXCL-10 production, directly correlating with increased disease severity and mortality. This establishes the cellular source of the pathogenic inflammatory signals identified in our cytokine analysis. Conversely, *Tg* infection promotes a neuroprotective A2 phenotype distinguished by enhanced TGF-β secretion, providing the cellular mechanism for anti-inflammatory cytokine production. In our study, we observed an increase in the number of astrocytes and microglia in the brains of mice coinfected with *Pb*A and *Tg* compared to uninfected controls. Interestingly, the abundance of these glial cells in coinfected mice was lower than in mice mono-infected with *Tg*, correlating with the reduced number of *Tg* cysts observed in the brains of coinfected mice. This suggests that *Pb*A coinfection influences *Tg* replication, potentially through modulation of the local immune environment. Critically, *Tg*-*Pb*A coinfected mice displayed astrocytes exhibiting an intermediate A1/A2 phenotype, maintaining balanced pro-inflammatory CXCL-10 and anti-inflammatory TGF-β production. Furthermore, reduced activation of glial cells in coinfected brains, characterized by an intermediate M1/M2 microglial activation profile. This reduced glial activity may reflect a more balanced neuroinflammatory response, which could simultaneously limit *Tg* replication while preventing excessive inflammation that contributes to cerebral malaria pathology. These findings suggest that the moderated glial response in coinfected mice may play a dual protective role: controlling *Tg* cyst formation and mitigating the neuropathological consequences of *Pb*A infection. This novel transitional activation state represents the cellular mechanism underlying the balanced cytokine profile and provides the direct link between parasitic co-presence and neuroprotective outcomes. The functional plasticity of astrocyte populations demonstrates how coinfection creates qualitatively distinct cellular responses rather than simply quantitative modifications of single-pathogen responses.

Building upon the astrocyte phenotype switching mechanisms, we identified a specific regulatory circuit that sustains and amplifies the neuroprotective response. Activated astrocytes produce substantial quantities of IL-33, correlating with significant recruitment of type 2 innate lymphoid cells (ILC2s) within the cerebral parenchyma during both *Tg* monoinfection and coinfection scenarios. IL-33 functions as a critical mediator of neuroimmune interactions, released as a damage-associated molecular pattern (DAMP) following cellular injury and promoting ILC2 recruitment and activation through its cognate receptor ST2 (31,46). This establishes a mechanistic pathway from initial tissue damage to protective immune cell recruitment. The IL-33/ST2 signaling axis promotes expansion of IFN-γ^+^ T lymphocytes, iNOS^+^ monocytes, and ILC2s, collectively contributing to *Tg* containment and cerebral defense against parasitic invasion(31). Recruited ILC2s subsequently produce IL-13 and IL-5, driving type 2 immune responses that provide tissue protection and neuroinflammation modulation (47). This creates a self-reinforcing protective cascade initiated by astrocyte activation. The interaction between IL-33 and ST2 receptors on astrocytes creates a novel autocrine regulatory circuit that sustains cytokine production while modulating the overall CNS immune environment during polymicrobial infections. Our data demonstrate that IL-33, through ST2 receptor engagement on astrocytes, maintains autocrine cytokine production in *Tg*-*Pb*A coinfected neural tissue, creating a self-perpetuating protective loop. This self-sustaining mechanism ensures that protective responses persist throughout the duration of coinfection, preventing the typical progression to fatal ECM. The identification of differential IL-33 gene variant expression patterns, with IL-33a correlating with enhanced survival and IL-33b mediating *Pb*A resistance, suggests that multiple parallel protective pathways operate simultaneously through variant-specific mechanisms.

Our observations align with extensive clinical evidence demonstrating that coinfections substantially modulate immune responses and disease outcomes, particularly in endemic regions where multiple pathogens exhibit co-transmission patterns (1–3,48,49). The mechanistic insights from our model provide a framework for understanding these clinical observations and suggest that apparent disease resistance in certain populations may reflect underlying coinfection dynamics rather than genetic resistance alone. The *Tg*-*Pb*A coinfection model provides a valuable experimental paradigm for investigating CNS disorders characterized by neuroinflammation and BBB dysfunction, with direct implications for understanding diseases such as multiple sclerosis and Alzheimer’s disease. The identified regulatory circuits may represent conserved protective mechanisms that could be therapeutically activated in these conditions. The mechanistic pathway we have elucidated—from parasitic co-localization through astrocyte phenotype switching to IL-33/ST2/ILC2 circuit activation—identifies multiple potential therapeutic intervention points. The IL-33/ST2 pathway represents a particularly promising therapeutic target for diseases characterized by severe neuroinflammation, including CM. Therapeutic modulation of this pathway may provide novel strategies for disease progression control in CNS-related disorders (50). By harnessing these endogenous immune regulatory mechanisms, novel treatment modalities may be developed to attenuate neuroinflammation, enhance disease tolerance, and effectively address the complexities of polymicrobial infections affecting the CNS.

Our findings establish a mechanistic framework wherein chronic *Tg* infection creates the necessary conditions for neuroprotection through sequential molecular and cellular events: spatial co-localization requirements, cytokine network remodeling, astrocyte phenotype switching, and self-perpetuating IL-33/ST2/ILC2 regulatory circuits. This progression from prerequisite conditions through specific molecular mechanisms to sustained protective responses provides a comprehensive understanding of coinfection-mediated neuroprotection and identifies multiple therapeutic targets for clinical intervention. The critical importance of incorporating coinfection dynamics into disease management paradigms extends beyond malaria to encompass broader neuroimmune disorders where similar protective circuits may be therapeutically activated.

## Data availability

Data will be made available on request.

## Acknowledgments

The authors acknowledge use of the PLETHA platform of Pasteur Institute of Lille for assistance in animal care and maintenance and are thankful to Dr. Hélène Bauderlique and Dr Olivier Molendi-Coste for technical assistance in flow cytometry and Dr Sophie Salomé-Desnoulez in confocal microscopy (BiCel Platform; Institut Pasteur de Lille). We also thank Prof Pierre-André Cazenave and Dr Jacques Roland (CIIL, Institut Pasteur and Sorbonne University) for critical evaluation of the manuscript and Mr David Fraser, Biotech communication for content editing.

This work was supported by the LabEx PARAFRAP: ANR-11-LABX-0024 for IL salaries.

## Author contributions

SP, SM: Study concept an design. IL, TK, FH, CP, MA: Performed the experiments. IL : statistical analysis. IL, TK, FH, SM: Contributed to reagents/materials/analysis tools. IL, TK: acquisition and interpretation of data. IL, TK, CG, SM, SP: contribute to writing of the manuscript.

## Supplementary data

**Supplementary figure 1.**
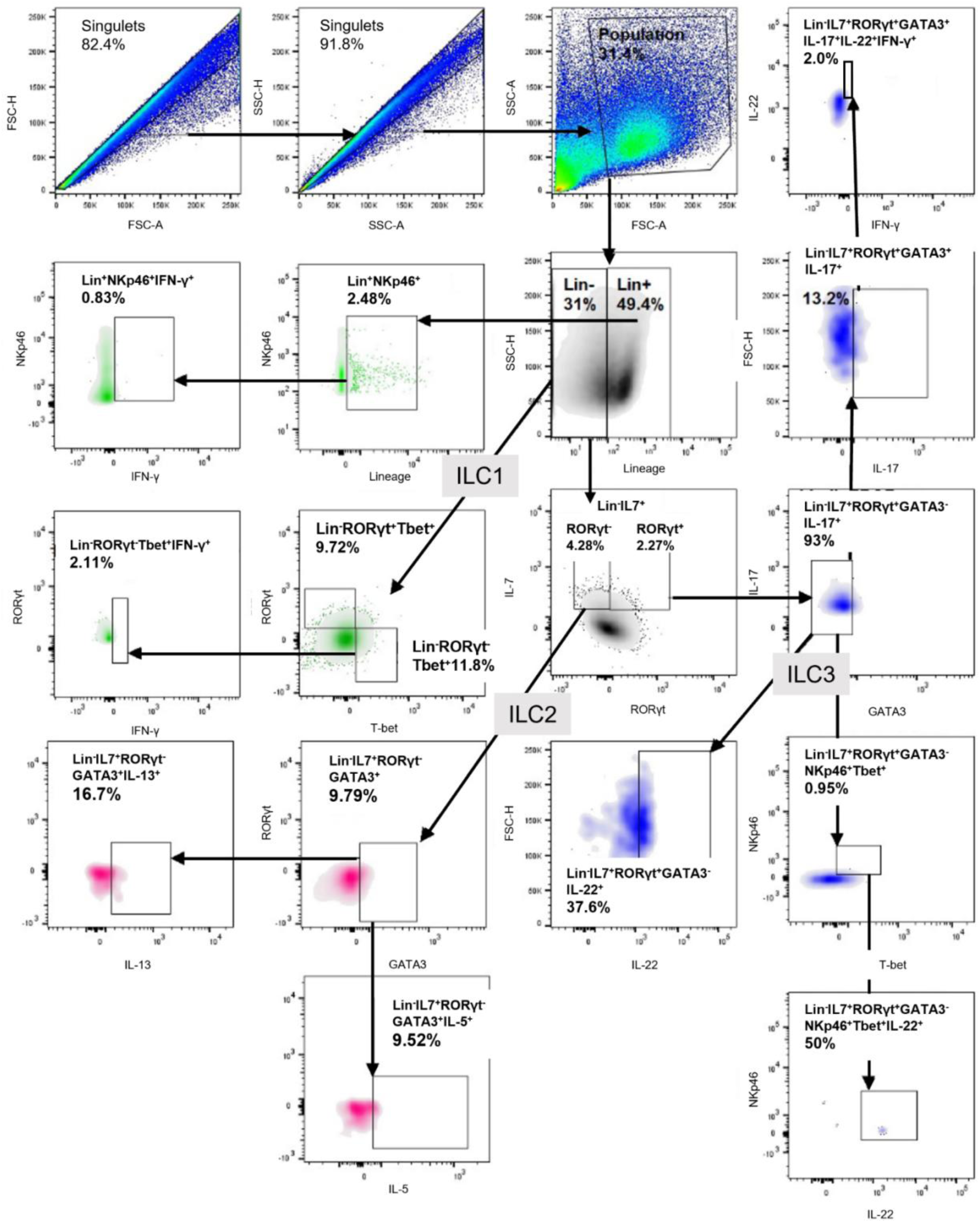
Gating strategy to target ILCs types by flow cytometry. ILC1 were labelled by Lineage^-^ RORγt^+^Tbet^+^; ILC2 by Lineage^-^IL-7^+^RORγt^-^GATA3^+^; and ILC3 by Lineage^-^IL-7^+^RORγt^+^GATA3^-^ IL-22^+^/NKp46^+^Tbet^+^.

**Supplementary Table 1.**
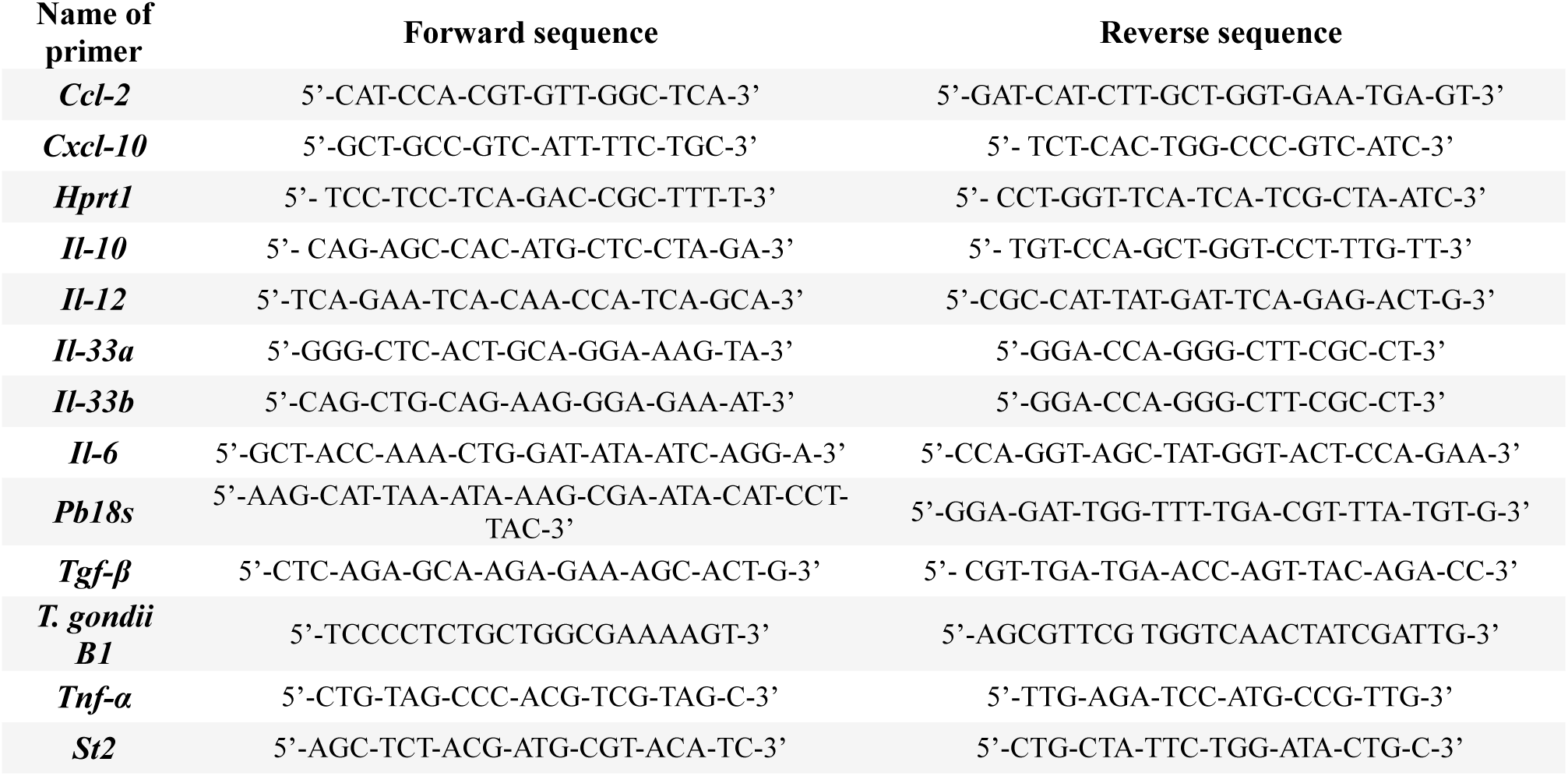
List of primers used for RT-qPCR analysis.

**Supplementary Table 2.** List of antibodies used in flow cytometry.

| Target | Fluorochrome | Source | Reference | Dilution |
| --- | --- | --- | --- | --- |
| **CCR2** | FITC | Biolegend | 150608 | 1/100 |
| **CD11b** | Pacific blue | Biolegend | 101224 | 1/400 |
| **CD16/32** | APC/Cy7 | Biolegend | 101328 | 1/100 |
| **CD206** | PerCP/Cy5.5 | Biolegend | 141715 | 1/1800 |
| **CD45** | PE/Cy7 | Biolegend | 103114 | 1/900 |
| **CD62L** | PerCP | Biolegend | 104429 | 1/50 |
| **CD86** | BV605 | Biolegend | 105037 | 1/200 |
| **CX3CR1** | BV605 | Biolegend | 149027 | 1/50 |
| **CXCL-10** | AF555 | Bioss | BOSSBS-150R-A555 | 1/300 |
| **GATA3** | AF647 | BD Pharmingen | 560068 | 1/50 |
| **GATA3** | FITC | eBioscience | 53-9966-42 | 1/100 |
| **GFAP** | AF488 | Invitrogen | 53-9892-82 | 1/100 |
| **GLAST** | APC | Miltenyi Biotec | 130-095-814 | 1/25 |
| **IFN-γ** | BV510 | Biolegend | 505841 | 1/50 |
| **IL-5** | PE | Biolegend | 504303 | 1/200 |
| **IL-13** | PE-eFluor610 | eBioscience | 61-7133-80 | 1/100 |
| **IL-17A** | APC/Cy7 | Biolegend | 506939 | 1/200 |
| **IL-22** | PerCP/Cy5.5 | Biolegend | 516411 | 1/100 |
| **IL-33** | PE | Biotechne | IC3626P | 1/50 |
| **CD127 (IL-7Rα)** | PE/Cy5 | Biolegend | 135013 | 1/400 |
| **Lineage** | Pacific Blue | Biolegend | 133310 | 1/25 |
| **Ly6C** | BV510 | Biolegend | 128033 | 1/100 |
| **Ly6G** | APC | Merck | MABF469 | 1/200 |
| **MHC class I (H2-K<sup>d</sup>)** | AF700 | Biolegend | 116627 | 1/300 |
| **MHC class II (I-A/I-E)** | PerCP/Cy5.5 | Biolegend | 107625 | 1/400 |
| **NKp46** | Pe/Cy7 | Biolegend | 137617 | 1/200 |
| **RORγt** | APC | eBioscience | 17-6981-82 | 1/200 |
| **RORγt** | BV650 | BD Pharmingen | 564722 | 1/50 |
| **ST2** | BV785 | Biolegend | 145321 | 1/50 |
| **Streptavidine** | BV711 | Biolegend | 405241 | 1/100 |
| **Tbet** | BV711 | Biolegend | 644820 | 1/200 |
| **TGF-β** | Biotine | Novus-eBiotechne | NBP2-45137B | 1/100 |

